# A novel multidimensional reinforcement task in mice elucidates sex-specific behavioral strategies

**DOI:** 10.1101/690750

**Authors:** Munir Gunes Kutlu, Jennifer E Zachry, Lillian J Brady, Patrick R Melugin, Christina Sanders, Jennifer Tat, Amy R Johnson, Kimberly Thibeault, Alberto J. Lopez, Erin S. Calipari

## Abstract

**Background:** Sex is a critical biological variable in the neuropathology of psychiatric disease, and in many cases, women represent a vulnerable population. It has been hypothesized that sex differences in neuropsychiatric disorders are manifestations of differences in basic reward processing. However, preclinical models often present rewards in isolation, ignoring that ethologically, reward seeking requires the consideration of potential aversive outcomes.

**Methods:** We developed a Multidimensional Cue Outcome Action Task (MCOAT) to dissociate motivated action from cue learning and valence. Mice are trained in a series of operant tasks. In phase 1, mice acquire positive and negative reinforcement in the presence of discrete discriminative stimuli. In phase 2, both discriminative stimuli are presented concurrently allowing us to parse innate behavioral strategies based on reward seeking and shock avoidance. Phase 3 is punished responding where a discriminative stimulus predicts that nose-poking for sucrose occurs concurrently with footshock, allowing for the assessment of how positive and negative outcomes are relatively valued.

**Results:** Females prioritize avoidance of negative outcomes over seeking positive, while males have the opposite strategy. In cases where rules are uncertain, males and females employ different strategies, with females demonstrating bias for shock avoidance.

**Conclusions:** The MCOAT has broad utility for neuroscience research where pairing this task with recording and manipulation techniques will allow for the definition of the discrete information encoded within cellular populations. Ultimately, we show that making conclusions from unidimensional data leads to inaccurate generalizations about sex-specific behaviors that do not accurately represent ground truth.

## Introduction

Most of our understanding about behavioral strategies gained from preclinical research relies on unitary measures of behavior where animals have one option in the environment (take drug or not, avoid shock or not) and often across only one reinforcer minimizing relative valuation of information. As such, unidimensional behavioral tasks are not sufficient to gain a holistic understanding of behavioral functions, or how they may manifest themselves in pathologies. In recent years, research focused on understanding the biological variables contributing to psychiatric disorders has highlighted sex-based differences in the development and presentation of symptoms as well as in fundamental behavioral processes (1–4). Understanding the factors contributing to sex-specific vulnerability to neuropsychiatric disease is critical to developing treatments that are safe and effective for both sexes. Thus, developing animal models that are capable of assessing complex behavioral strategies while remaining quantitative and readily interpretable is critical for gaining understanding of the interplay between behavioral strategies and neuropsychiatric disease through preclinical research.

A great deal of previous research has highlighted that assessing how animals weigh environmental information to guide behavioral strategies can be highly complex, both within and between sexes. For example, even though females will self-administer opiates at higher rates than males (2), when given a choice between opiates and a high-fat reward they choose the non-drug reinforcer over the drug alternative (5), clearly highlighting that sex-differences do not manifest themselves as universal behavioral principles, but rather are a complex interaction between sex and environment. Capturing this complexity necessitates behavioral tasks that can probe the balance in the subjective value of rewarding versus aversive stimuli - and their antecedent cues - and how this balance, or bias, can be shifted in altered states - either biological or otherwise. Together making conclusions from unidimensional data can lead to broad scale generalizations about sex-specific behaviors that do not accurately reflect sex differences in behavioral strategies.

At the center of psychiatric disorders, such as substance use disorder, anxiety, and depression is a dysregulation in reward and motivation (6). In practical contexts, rewards are generally weighed with potential negative consequences (7), requiring consideration of the value of a reward and of associated aversive outcomes. Sex-based differences in reward seeking and avoidance - developed over evolutionary history - offer an ideal model to explore bias and strategy in a behavioral task while providing insight into the neurobiological basis of information encoding (8–12). Previous work has shown that the schedule under which a stimulus is presented is more of a behavioral determinant than the stimulus itself (13, 14); while these principles laid the ground-work for behavioral neuroscience, many of these fundamental findings are no longer considered when developing and designing behavioral models in animals.

Here, we establish and validate a novel rodent behavioral task that allows for quantitative assessment of multidimensional behavioral functions relevant to human decision-making. In this task, deemed Multidimensional Cue Outcome Action Task (MCOAT), mice are trained to respond to discriminative stimuli that predict either positive (nose-poke to get sucrose) or negative (nose-poke to avoid shock) reinforcement, then subsequently presenting both cues concurrently in conflict trials. In this task, we can probe behavioral strategies, and can also examine the neurological basis of mechanisms that subserve decisions regarding the subjective value of a reward and how the balance between rewarding and aversive stimuli can shift with training and in contexts of limited or complete information. We apply this approach in male and female mice to demonstrate latent sex-specific behavioral strategies that are exposed at times of conflict or uncertainty in optimal behavioral strategies, which would not be apparent using behavioral tasks where a single option is presented. We show in this study a significant sex-based difference in the response to positive versus negative reward-predictive cues, where females demonstrated a partiality to avoid shocks while males did not exhibit a bias toward any particular outcome. Additionally, this disparity between male and female mice dissipates as the level of familiarity with the available outcomes is increased. Together these data highlight fundamental sex-specific behavioral strategies guiding goal-directed behavior and highlight the importance of multidimensional tasks in understanding behavioral control.

## Methods and Materials

### Animals

Male and female 6-to 8-week-old C57BL/6J mice were housed five per cage. Mice were obtained from Jackson Laboratories (Bar Harbor, ME; SN: 000664). All animals were maintained on a 12h reverse light/dark cycle. Animals had free access to water but were food restricted to 90% of free-feeding weight for the duration of the studies. Mice were weighed every other day to ensure that weight was maintained. Animals were fed 2.5g chow per/mouse/day and food intake was adjusted to meet the weight criteria based on animals body weight each day. Behavior was conducted during the dark phase of the light cycle. All experiments were conducted in accordance with the guidelines of the Institutional Animal Care and Use Committee at Vanderbilt University School of Medicine, which approved and supervised all animal protocols. Experimenters were blind to experimental groups and the order of testing was counterbalanced during behavioral experiments.

### Apparatus

Mice were trained and tested daily in individual Med Associates (St. Albans, Vermont) operant conditioning chambers fitted with a house light, grid floor with shock harness, programmable tone generator, speakers, and two illuminated nose-pokes on either side of a sucrose delivery port equipped with an infrared beam break to assess head entries. One nose-poke functioned as the active operanda, and the other as the inactive, depending on the phase of the experiment (described below). Responses on both nose-pokes and head entries into the sucrose port were recorded throughout the duration of the experiments.

### Multidimensional cue outcome action task (MCOAT)

*Experimental Timeline*: Animals were trained in a series of operant tasks, for which they must meet task-specific criteria (defined below) before moving to the next phase. Mice that fail to meet each criterion do not continue to the next phase - the percentage of animals in each group completing criteria is outlined in Figure 2 and Supplementary Fig 3. The different phases of the task are (***Phase 1***) positive reinforcement and negative reinforcement, (***Phase 2a***) limited discrimination and conflict, (***Phase 2b***) extensive discrimination and conflict, (***Phase 4***) and punished responding (Fig 1). Animals were trained in one 1h session daily. All experiments were done within subjects allowing for comparisons to be made across training sessions and conditions.

**Figure 1.**
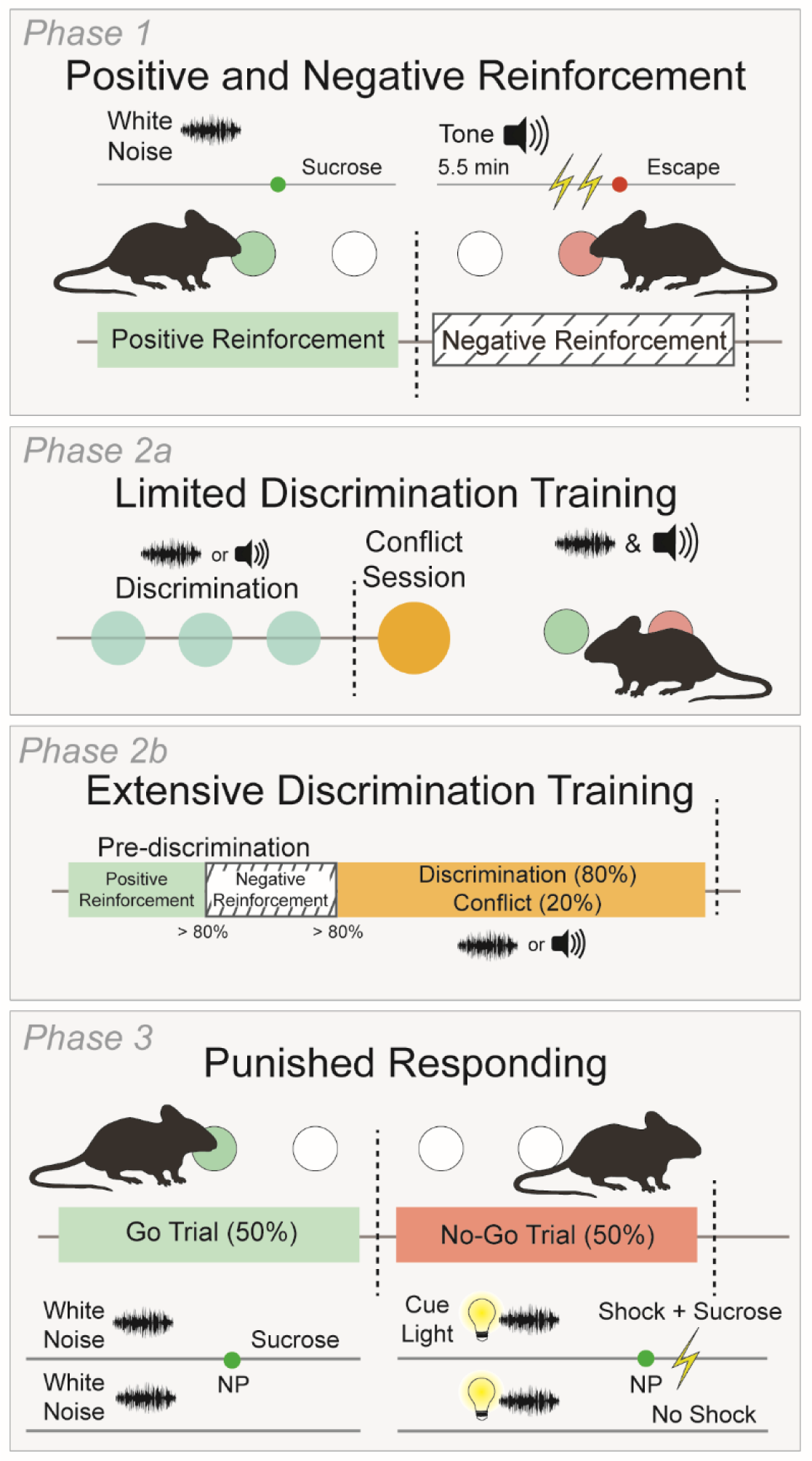
Schematic of the Multidimensional Cue Outcome Action Task (MCOAT). In Phase 1, of the MCOAT mice are first trained in positive reinforcement on an FR1 schedule. A discriminative stimulus (Sd1, white noise or tone counterbalanced between animals and conditions) was presented for the duration of the session indicating that nose-pokes on a defined side (either left or right) were reinforced by sucrose delivery. In the second component of Phase 1, mice acquired negative reinforcement. In these trial-based sessions a separate auditory Sd (Sd2) was presented to indicate that nose-poking (on the opposite nose-poke) was reinforced by the removal of a series of foot shocks. Following acquisition mice transitioned to Phase 2a which had a discrimination phase (80% of trials), where each Sd is randomly presented, and animals are required to emit the correct operant response: poke for sucrose or poke for shock removal - depending on the Sd presented. In the remaining trials (20%), Sd1 and Sd2 are presented simultaneously (Sd1+2) and animals have the option to nose-poke to obtain a sucrose reward or nose-poke to avoid shock. Thus, intrinsic response biases can be assessed. In Phase 2b, mice are trained until they meet a discrimination criterion of >70% and thus, have extensive experience with the discrimination/conflict portion of the task to understand how response bias changes with training experience. Finally, in Phase 3, animals are trained that a compound cue predicts punishment. Briefly, in 50% of the trials Sd1 is presented and predicts positive reinforcement. In the remaining 50% Sd1 is presented with a secondary Sd3 (Sd1+3) that predicts that a nose-poke will result in the delivery of sucrose and a footshock simultaneously. Together this behavioral task allows for the assessment across a wide range of approaches within individual animals.

**Figure 2.**
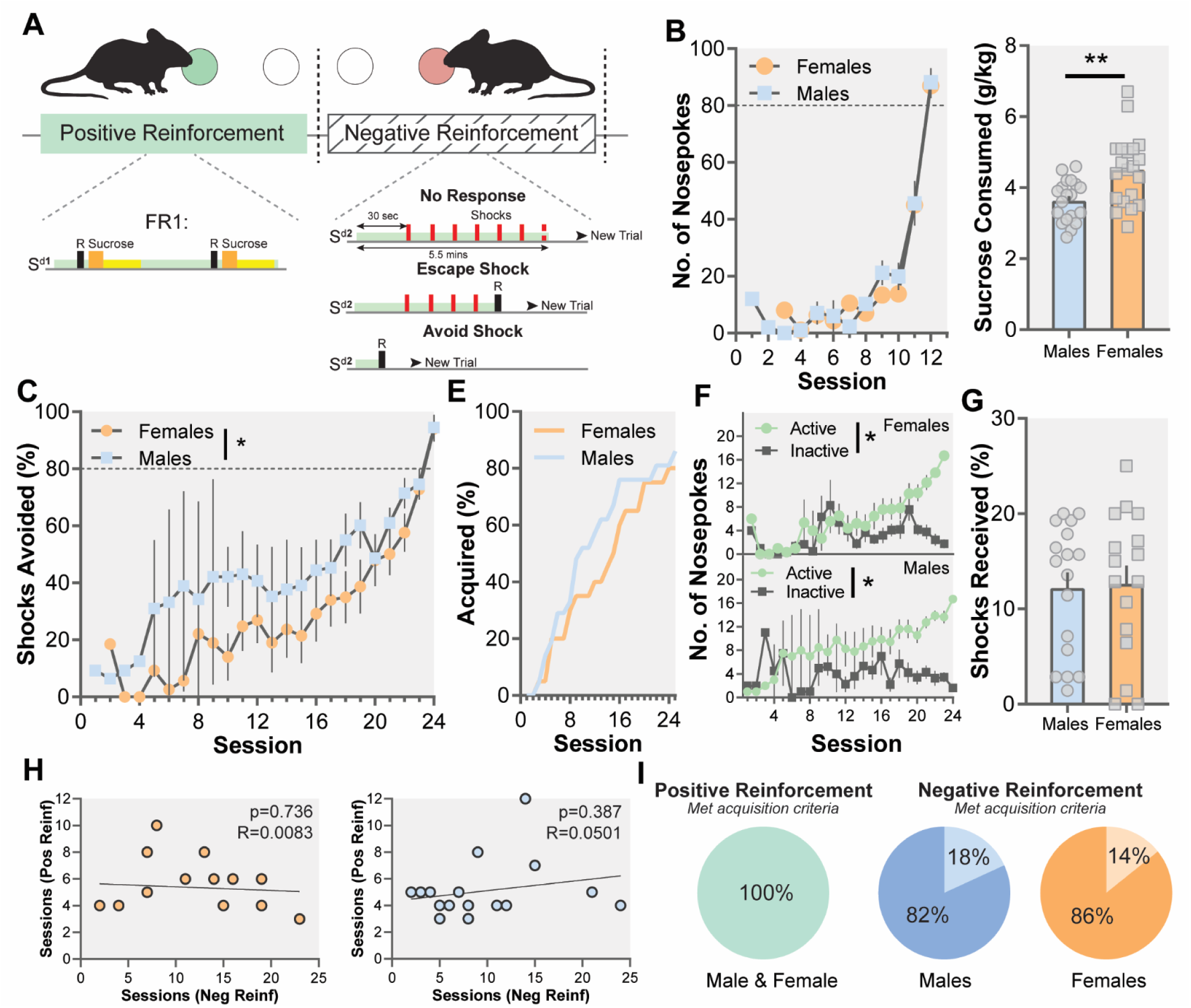
Sex-differences in reinforcement for positive and negative reinforcement. (A) Schematic of reinforcement schedules. Mice were trained on positive reinforcement and negative reinforcement. (B) Males and females learned positive reinforcement at similar rates. (C) Females consume more sucrose than males (t(38)= 2.603, p=0.0131). (D) Males acquired negative reinforcement at a faster rate than females (Mann-Whitney U= 164, p=0.0164). (E) Survival curve showing sex differences in the acquisition of negative reinforcement. (F) Both males (Mann-Whitney U= 96.50, p<0.0001) and females (Mann-Whitney U= 150.5, p=0.0114) showed significant differences in the number of active and inactive nose-pokes. Males and females mice that acquire the task did not differ in performance showing that the effect is selective to acquisition (X2 (22) = 14.97, p>0.05). (G) There was no difference in the number of shocks received once males and females reached criterion (t(30), 0.5210, p>0.05). (H) There is no relationship between the number of sessions to criterion for positive and negative reinforcement sessions in males or females showing that the task phases are independent measures. (I) Pie charts showing the percentage of animals that completed each component of phase I of the MCOAT. Data represented as mean ± S.E.M. * p < 0.05, ** p < 0.01, *** p < 0.0001.

#### Phase 1: Positive and Negative Reinforcement

##### Positive Reinforcement

Mice were trained on a fixed-ratio 1 (FR1) schedule of reinforcement to nose-poke in the active poke for sucrose delivery (1s duration of delivery, 10uL volume, 1mg sucrose). Upon each correct response the sucrose delivery port was illuminated for 5 seconds and sucrose was delivered. During *Phase 1* training sessions, an auditory discriminative stimulus (**S**^**d1**^) - white noise or 2.5 khz tone (counterbalanced) - was presented for the entirety of the session. Mice were moved to the next phase when they responded on the active NP >80 times in a session.

##### Negative Reinforcement

Mice were trained to nose-poke on the opposite, non-sucrose-paired nose-poke for negative reinforcement - to prevent the presentation of foot shocks. All shocks were short, but high intensity: 1.0 mA in magnitude delivered for 0.5s. A second auditory discriminative stimulus (**S**^**d2**^) -- either tone or white noise, counterbalanced between positive and negative reinforcement -- was presented on a variable interval 30s (VI30) schedule for the inter-trial interval (ITI). At the beginning of each trial, **S**^**d2**^ came on for 30s after which a series of shocks was delivered (15 second inter-stimulus interval (ISI), 20 shocks total). In this task, mice are able to respond any time during the trial to end the trial and begin the ITI. Responding on the correct nose-poke during **S**^**d2**^ immediately ended the trial, thus preventing the shocks from being presented (avoidance). If responses were made after shocks commenced, responding on the correct nose-poke terminated the shocks and ended the trial (escape). The shock and **S**^**d2**^ were terminated immediately following a correct response. The trial ended either after the animal made a correct response or after 330 seconds. Acquisition criteria was defined as receiving fewer than 25% of total possible shocks in a session. Animals that did not meet this criterion after 15 sessions were removed from the study.

#### Phase 2a: Limited Discrimination and Conflict

Following acquisition of both the positive and negative reinforcement tasks, mice went into *Phase 2a*. In the limited discrimination phase, mice were trained in one 1h session per day for three consecutive days. In this trial-based phase, 80% of the trials were discrimination trials and 20% were conflict trials:

##### Limited Discrimination Pre-Training

Before the beginning of the conflict testing, animals underwent three sessions of discrimination only training to ensure that they were using the antecedent cues (**S**^**d1 OR 2**^) to guide their operant responses. In these trials, **S**^**d1**^ and **S**^**d2**^ were presented in random order and equal proportion and responses on the correct (corresponding to the **S**^**d**^ that was presented) and incorrect nose-poke were recorded. The **S**^**d**^ predicted the same response between phase 1 and 2, the only difference is that they were presented randomly within the same session to ensure that the animals had acquired the association. Depending on the S^d^, animals could respond on the appropriate nose-poke for either sucrose or shock avoidance. Response on the active nose-poke during **S**^**d1**^ initiated a 1s sucrose delivery and terminated the **S**^**d1**^, effectively ending the trial. Response at the opposing nose-poke during **S**^**d2**^ terminated **S**^**d2**^ and ended the trial. Failure to make an active response during the 30 second duration of the **S**^**d2**^ resulted in a single shock and the trial end. Mice that did not respond in either sucrose or shock trials were removed during this phase.

##### Discrimination and Conflict

The test session consisted of both discrimination trials (80% of trials) and conflict trials (20% of trials) in the same session. Discrimination trials were identical to those described above. In conflict trials, mice were presented a compound cue (**S**^**d1**^ + **S**^**d2**^) for 30 seconds. Both nose-pokes were illuminated. Depending on their response, mice received one of three possible outcomes: 1) failure to respond resulted in a shock at the end of the 30s compound cue, 2) if they responded on the sucrose active side, they received sucrose and a footshock, 3) If they responded on the negative reinforcement active side, they avoided shock and did not receive sucrose. As before, trials and **S**^**d**^s were terminated following an active response. This allowed us to define animals’ response bias when conflicting information was presented.

#### Phase 2b: Extensive Discrimination and Conflict

Following acquisition of both the positive and negative reinforcement tasks, a second cohort of mice underwent extensive discrimination training. Each day, mice underwent a 15-minute pre-discrimination positive reinforcement session, a 15 min pre-discrimination negative reinforcement session, and a 1hr discrimination/conflict session (80% discrimination trials, 20% conflict trials as described above). In both the positive and negative reinforcement sessions, the mice responded in >80% of the trials to move onto the next session for that day. Mice were trained daily in discrimination until they reached a criterion of >70% correct.

#### Phase 3: Punished Responding

Mice trained in positive and negative reinforcement and that underwent limited discrimination and conflict (*Phase 2*) were moved to punished responding. Each session contained 50% positive reinforcement trials for sucrose and 50% punished trials.

##### Positive reinforcement trials

Mice were presented **S**^**d1**^ and had 30 seconds to nose-poke on the active poke for sucrose. Sucrose was delivered as described above. **S**^**d**1^ and the trial were terminated following an active response or at the end of the 30 seconds.

##### Punished trials

**S**^**d1**^ and **S**^**d3**^ (a house light) were presented concurrently. Responding on the active nose-poke on these trials resulted in the delivery of sucrose and a single footshock. The intensity of this shock was increased in each subsequent session over the course of 9 total sessions (0.1, 0.2, 0.3, 0.4, 0.5, 0.6, 0.75, 1.0, and 1.5 mA). In these trials to avoid shock the animal must inhibit behavioral responding - thus the outcome is the same as in phase 1 (avoid shock) but the behavior is opposite (go vs no go).

### Shock Sensitivity

To rule out differences in shock sensitivity between males and females as a factor contributing to the behavioral outcomes, a follow-up shock sensitivity task was conducted after the punished responding phase. During one 1h session, animals received randomly selected magnitude shocks of 0.1, 0.2, 0.3, 0.4, 0.5, 0.6, 0.75, 1.0, or 1.5 mA with variable ITI of 30, 45, or 60 seconds. All shocks were unsignaled and no cues were presented in the session. Vocalization (non-ultrasonic) and motor responses were scored. Vocalization was scored as a 1 if the subject vocalized and a 0 if the subject did not vocalize in the session. Motor responses were scored as a 1 if the subject ran, a 2 if the subject hopped (4 paws off the ground), a 3 if the subject ran and hopped, and as a 0 if the subject did not move.

### Analysis Parameters

For positive and negative reinforcement tasks, the total sucrose and total shock responses were analyzed using unpaired t tests. Mann-Whitney U test was used when the number of sessions to the criterion was not equal between subjects precluding the use of parametric statistics. Discrimination and conflict task responses were analyzed using two-way ANOVA (Trial Type x Sex) when they were represented as averages or when they represent a single session (Limited Discrimination Phase). We employed a mixed Repeated Measures ANOVA for the Punished Responding and Shock Sensitivity experiments where all mice received the same number of sessions. We also used a computational analysis to determine the parameters of response bias (Log *b*) and discrimination (Log *d*), as described previously (15, 16). Briefly, Log *d* value was derived mathematically as a measure of the rate of discrimination in a bias-independent measure whereas Log *b* was computed as the measure for behavioral bias. Both terms use a logarithmic scale for the multiplication of the ratio between correct and incorrect responses during two different trial types. We explain the mathematical terms in detail below:

### Log d

*Log d* is a measure of the rate of discrimination for the S^d^. In the discrimination phase of the MCOAT, mice were trained to nose-poke in the right or left poke based on the S^d^ presented to them in order to obtain sucrose or avoid footshocks. Log *d* is determined as the ratio between the number of correct and incorrect Sucrose and Shock trials, which results in a negative (no discrimination) or a positive (successful discrimination):

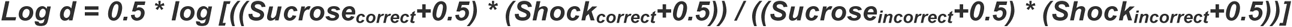

### Log b

Log *b* is a measure of the animals’ response bias. In our task, the animals were presented a compound stimulus consisting of two auditory cues signaling opposite outcomes. In these conflict trials, the subjects had to choose between getting a sucrose reward versus avoiding a footshock. Using the data from these trials, we assessed “behavioral bias,” that is, the preference of mice for one outcome over another. Log *b* is calculated as the ratio between the number of correct Sucrose and incorrect Shock versus incorrect Sucrose and correct Shock trials, which results in either a negative (bias towards avoidance) or a positive value (bias towards sucrose) or a 0 (no bias):

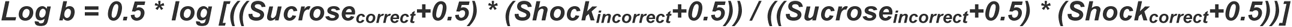

## Results

### Females show increased measures of positive reinforcement and a decreased learning rate for negative reinforcement

In the MCOAT, *Phase 1* requires mice to acquire both positive and negative reinforcement. Positive and negative reinforcement have the same behavioral response (nose-poke) and both have positive outcomes (sucrose delivery, removal of a negative stimulus) but the valence of the stimulus maintaining the responding is divergent (shock vs sucrose). We can thus examine how male and female mice learn positive outcomes with different reinforcers (negative vs. positive; Fig 2A). Our results showed that although females consumed more sucrose during each reinforcement trial (Fig 2B; *t*(38)= 2.603, *p*=0.0131), there was no significant difference in the positive reinforcement learning rate in males and females (Fig 2B; *p* > 0.05). In contrast, we found that males performed better in the negative reinforcement task compared to female mice and avoided a significantly larger percentage of shocks during the acquisition phase (Fig 2C; Mann-Whitney U= 164, *p*=0.0164). However, both males (Mann-Whitney U= 96.50, *p*<0.0001) and females (Mann-Whitney U= 150.5, *p*=0.0114) showed significant differences in the number of active and inactive nose-pokes (Fig 2F), demonstrating that both groups successfully learned the task. In addition, the percentage of male and female mice that completed the task did not differ (Fig 2E,J; *X*^*2*^ (22) = 14.97, *p*>0.05) and there was no difference in the number of shocks male and female mice received once they reached the criterion (Fig 2G; *t*(30), 0.5210, *p*>0.05). Because these tasks were all completed within subject, the effects must be stimulus-specific as females and males learned positive reinforcement at similar rates, ruling out differences in learning and memory capabilities between groups.

One of the major components of this task is to dissociate different behavioral strategies; however, for that to be possible each phase of the task must be independent from one another. Indeed, we found no significant correlation between the number of sessions to criterion for positive and negative reinforcement in males or females (Fig 2H; *p* > 0.05). Indicating that each aspect of the task is a dissociable component that tests different aspects of behavior. This also confirms that the sex differences that were observed in the negative reinforcement phase of the task are not a function of differences in overall learning rate between males and females.

### Females’ responses are biased towards avoiding negative outcomes when conflicting information is presented

In *Phase 2a* of the task, animals are presented with limited discrimination training and sessions where conflicting information is presented that requires them to make decisions about whether they value avoiding negative stimuli or seeking out positive. During the discrimination and conflict phases of our task, mice first received a limited number of discrimination training sessions to confirm that mice were in fact discriminating the **S**^**d**^s from one another, followed by a test session where 80% of the trials were discrimination trials and 20% were conflict trials (Fig 3A). During these trials, we calculated Log *d* to determine the level of discrimination for the **S**^**d**^ regardless of outcome and Log *b* which is a measure of response bias to define the behavioral bias during conflict trials. These computational measures of discrimination and bias are single measures that encompass the entirety of the bias and allow for correlations across other aspects of the task. Together, this design allowed for us to define sex-specific response biases that were not due to alterations in the ability of animals to discriminate the **S**^**d**^, but rather were intrinsic behavioral biases that guide decision-making.

**Figure 3.**
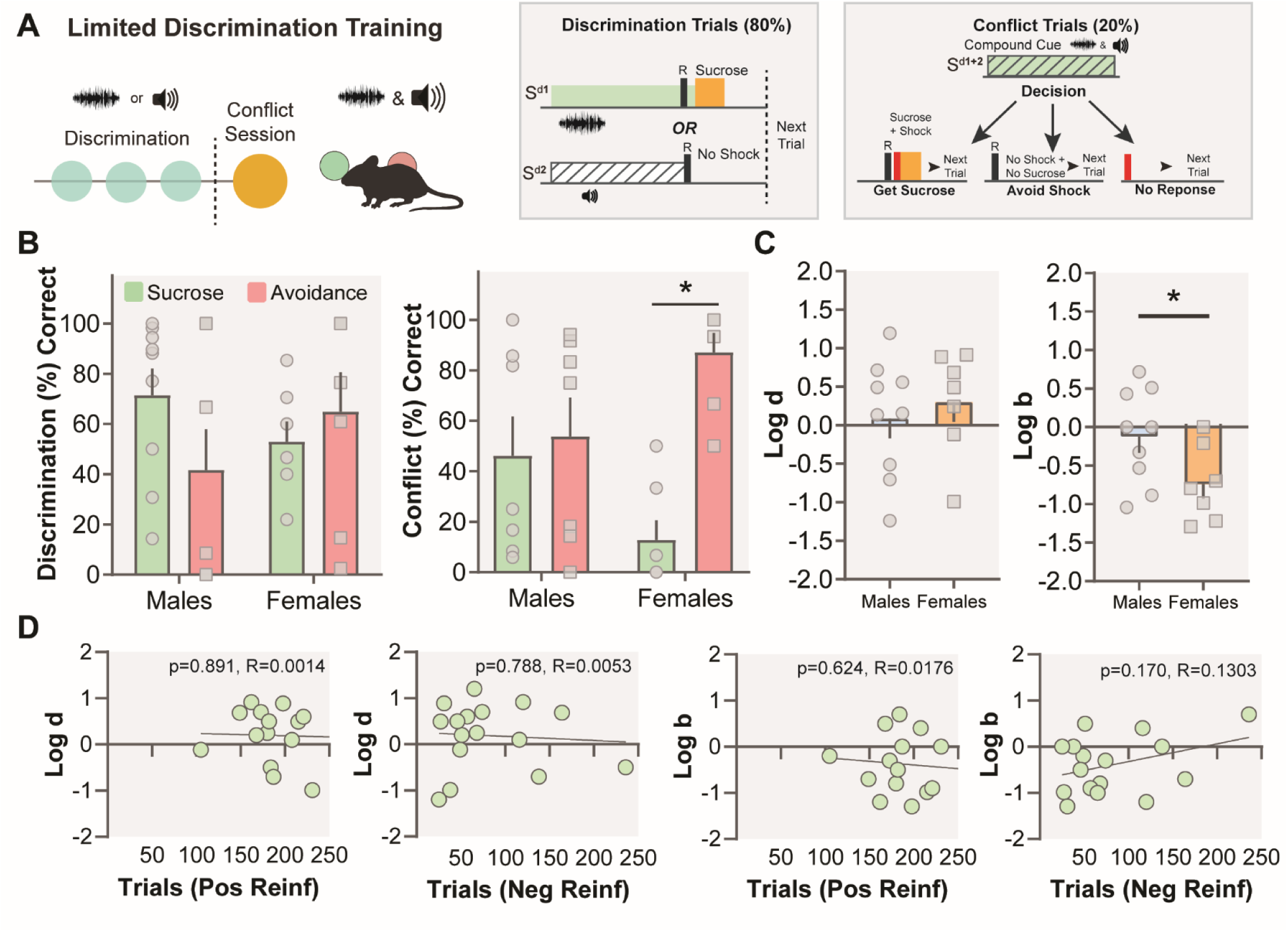
Females are bias towards shock avoidance. (A) In limited discrimination, mice are given three discrimination sessions. In the fourth session, 80% of the trials remain discrimination trials and 20% of the trials are conflict trials, in which both Sds are presented simultaneously. (B) Males and females show comparable levels of discrimination on the final discrimination session (Sex x Trial Type Interaction, F(1,14)=2.885, p>0.05; Sex main effect, F(1,14)=0.028, p>0.05). (B) In conflict trials, there is a significant interaction between Sex and Trial Type (Fig 3B F(1,24)=7.410, p=0.0119). The number of sucrose and avoidance responses was significant for females (t(6)= 4.816, p=0.0030) but not males (t(6)= 0.2448, p>0.05). (C) There is no significant main effect of Trial Type (F(1,14)=0.526, p>0.05) or sex on Log d (Fig 3C t(14), 0.5797, p>0.05). There is, however, a significant effect of sex on Log b (Fig 3C t(14)= 2.159, p=0.0487). Females show bias toward shock avoidance, while males do not demonstrate bias. (D) There is no statistically significant relationship between Log b and Log d values with the number of sessions to meet the criteria for positive or negative reinforcement.

Both sexes showed similar levels of discrimination, indicating that there were not sex differences in the ability of mice to discriminate antecedent cues (Fig 3B; Sex x Trial Type Interaction, F(1,14)=2.885, *p*>0.05; Sex main effect, F(1,14)=0.028, *p*>0.05) and Log *d* (Fig 3C; *t*(14), 0.5797, *p*>0.05). There was also no significant main effect of Trial Type (F(1,14)=0.526, *p*>0.05) indicating that animals completed similar numbers of sucrose and avoidance trials further dissociating any significant differences in bias from baseline responding.

During conflict trials, there was a significant interaction between Sex and Trial Type (Fig 2B; F(1,24)=7.410, *p*=0.0119) and a significant sex difference for Log *b* (Fig 3C; *t*(14)= 2.159, *p*=0.0487) demonstrating that male and female mice show differential biases towards sucrose and avoidance response. Specifically, while female mice chose to avoid shocks over sucrose, male mice did not bias for one of the competing responses. In line with these results, the difference between the number of sucrose and avoidance responses was significant for females (Fig 3B; *t*(6)= 4.816, *p*=0.0030) but not males (*t*(6)= 0.2448, *p*>0.05). Finally, we found no significant correlations between Log *b* and Log *d* values with the number of days to complete the positive or negative reinforcement task (Fig 3D; *p*>0.05), suggesting the learning rate in the initial tasks does not affect the performance or bias during the discrimination or conflict sessions. However, at the end of this phase of the task, only 50% of the females and 54% of the male subjects were able to complete the task with limited training due to the difficulty of the task for mice. This prompted us to go onto an extended discrimination phase with a separate group of mice (Supplementary Fig 3). Overall, these results clearly show that female mice show an intrinsic bias towards avoiding aversive outcomes over obtaining positive outcomes whereas male mice do not display the same bias.

### Females response bias does not change over extensive training, while males exhibit a bias towards shock avoidance following extensive training

In *Phase 2b* of the MCOAT, a series of additional training sessions to improve discrimination and familiarity with the task were conducted (i.e. Extended Discrimination Training and Conflict, Fig 4A). A second group of mice that completed *Phase 1* underwent the extensive discrimination training phase (Fig 4A). The goal of this phase was to examine: 1) the progression of discrimination learning and behavioral bias and 2) the behavioral bias during conflict when animals reach a set level of discrimination and are familiar with the task (>70% correct for both sucrose and shock trials). We found a significant interaction for Discrimination/Bias and Trial for males (Fig 4B; F(19, 154) = 3.366, *p*<0.0001) but not for females (Fig 4B; F(32, 98) = 1.219, *p*>0.05). In addition, there was a significant difference in the Log *d* (Mann-Whitney U= 221.5, *p*=0.0463) and Log *b* (Mann-Whitney U= 141, *p*=0.0003) values between males and females throughout the discrimination training. However, there was no significant interaction between Sex and Trial for Log *d* (F(4, 40) = 1.393, *p*>0.05) but the main effect of Trial was significant (F(2.684, 26.84) = 10.68, *p*=0.0001). There was also no difference in the number of days to criterion between sexes (Fig 4B; *t*(10)= 1.846, *p*>0.05). Furthermore, once male and female mice reached discrimination criterion they showed the same level of discrimination learning (Fig 4C; see Supplementary Fig 2 for correct, incorrect, and omission trial numbers and latencies; Sex x Trial Type Interaction, F(1,10)=0.1921, *p*>0.05). At the end of the discrimination training both sexes showed a significant response bias towards choosing avoidance over sucrose during the conflict trials (Fig 4C; Sex x Trial Type Interaction, F(1,20)=1.641, *p*>0.05; Trial Type Main Effect, F(1,20)=64.58, *p*<0.0001). Similarly, the Log *d* (*t*(10)= 0.9860, *p*>0.05) and Log *b* (*t*(10)= 0.1639, *p*>0.05) differences between sexes also disappeared following increased familiarity with the task structure (Fig 4D, E) suggesting that with extended training male mice show similar bias towards avoiding shocks over obtaining sucrose. Likely, because in the task omissions and sucrose responses both result in shock delivery, and the only way to avoid a shock is to respond on the poke paired with negative reinforcement. Giving further support to the idea that extensive experience with the task shifts the response bias, Log *b* and Log *d* values were negatively correlated for the limited discrimination phase (p<0.01, R=0.4794), indicating that the animals who showed better discrimination - and thus understood the task structure - were more inclined to avoid footshocks over obtaining sucrose (Supplementary Fig 1). Interestingly, while all male mice reached the discrimination criterion following extensive discrimination training, only 50% of the female mice successfully acquired discrimination (Supplementary Fig 3). These results highlight two important conclusions. First, with extensive discrimination training male mice developed a bias towards avoiding negative outcomes while female mice preserved their bias for shock avoidance regardless of experience or training. Second, female mice had a harder time learning the discrimination between sucrose and footshock trials.

**Figure 4.**
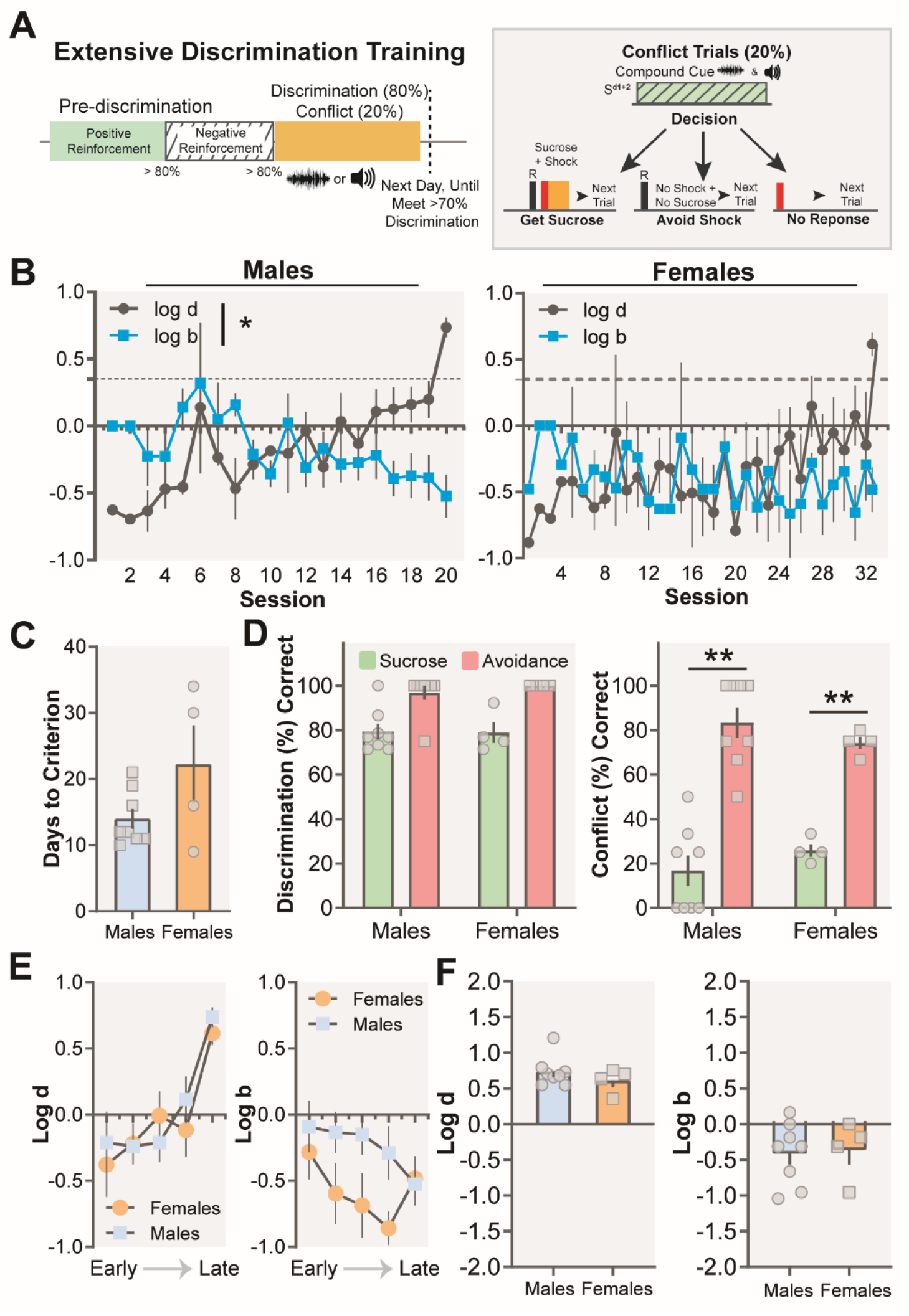
Extensive training on the MCOAT does not alter female bias towards avoiding aversive stimuli. (A) Schematic of extensive discrimination and conflict task. (B) In opposition to the findings with limited discrimination training, there was a significant interaction for Discrimination/Bias and Trial for males (F(19, 154) = 3.366, p<0.0001) but not for females (F(32, 98) = 1.219, p>0.05). (C) The difference in the number of days to criterion is not significant between sexes (t(10)= 1.846, p>0.05). (D) Once both groups met discrimination criterion, they demonstrated similar levels of discrimination (Sex x Trial Type Interaction, F(1,10)=0.1921, p>0.05). Following extended discrimination training, males and females both showed a response bias for shock avoidance in conflict trials (Sex x Trial Type Interaction, F(1,20)=1.641, p>0.05; Type Main Trial Effect, F(1,20)=64.58, p<0.0001). (E) There was no significant interaction between Sex and Trial for Log d (F(4, 40) = 1.393, p>0.05) but the main effect of Trial was significant (F(2.684, 26.84) = 10.68, p=0.0001). (F) There was no sex-differences in Log d (t(10)= 0.9860, p>0.05) and Log b (t(10)= 0.1639, p>0.05) following discrimination training.

### Female mice are more sensitive to aversive stimuli functioning as punishers

In the final phase of the MCOAT (*Phase 3*), mice are trained to self-administer sucrose either in the presence or absence of a cued punisher (a footshock paired with the nose-poke and delivery of sucrose). Punishers function to decrease rates of responding (17) and we tested the ability of male and female mice to reduce their rates of behavior in the presence of successive shocks of ascending intensity over sessions (Fig 5A). Our results showed that there was a significant difference between males and females in the number of sucrose responses during punishment (Fig 5C; Sex x Shock Intensity Interaction, F(8, 72) = 0.3219, *p*=0.9552; Sex Main Effect, F(1, 9) = 5.157, *p*=0.0493; Shock Intensity Main Effect, F(2.930, 26.37) = 15.16, *p*<0.0001) as well as for the number of shock responses (Fig 5C; Sex x Shock Intensity Interaction, F(8, 72) = 1.259, *p*>0.05; Sex Main Effect, F(1, 9) = 4.644, *p*=0.0596; Shock Intensity Main Effect, F(2.074, 18.67) = 41.02, *p*<0.0001) between varying shock intensities. In addition, the main effect of Trial Type is significant for males (Fig 5H; F(1, 10) = 10.90, p=0.0080) but not for females (Fig 5H; F(1, 8) = 3.812, p=0.0867) suggesting that both groups learned to differentiate between sucrose and punished trials; however, females were more sensitive to the effects of punishers on both trial types. We also computed response inhibition 50 curves (RI_50_) for each animal to determine the shock intensity value which caused a 50% reduction in behavioral responding (Fig 5D,F). There was a significant shift in the intensity required for the punisher to reduce behavior in males - i.e. females required lower shock intensities (were more sensitive to shock) - to reduce responding (Fig 5E; Mann-Whitney U=5, *p*=0.0480). This is particularly interesting, as the punisher is having greater effects in females, even on the trials where it is not presented (i.e. the unpunished trials). Female mice also show a tendency to learn to inhibit their responses for obtaining sucrose when an aversive outcome was signaled (Fig 5G; (*t*(9)= 1.971, *p*=0.1810). Overall, this is in line with our results showing female mice are biased to avoid negative outcomes, and this shows that this occurs in contexts where the avoidance is active (lever press to remove shock) or requires response inhibition (such as in punished trials). ***Females are less sensitive to unsignaled shocks***. Finally, it was important to assess whether differences in the unconditioned response to the shock could underlie these effects. To this end we measured vocalizations and motor responses to a variety of intensities of shocks. Our results showed that there was a significant sex difference in motor response (Fig 5B; Sex x Shock Intensity Interaction, F(4,35)=2.802, *p*=0.0406; Sex Main Effect, F(1,35)= 1.830, *p*>0.05; Shock Intensity Main Effect, F(4,35)= 48.33, *p*<0.0001) and in vocalization (Fig 5B; Sex x Shock Intensity Interaction, F(4,35)=6.991, *p*=0.0003; Sex Main Effect, F(1,35)= 27.58, *p*<0.0001; Shock Intensity Main Effect, F(4,35)= 111.3, *p*<0.0001) to different shock intensities. There is a sex difference in sensitivity to varying shock intensities. However, Bonferroni-corrected t-tests showed that only males showed higher motor response to 0.30 mA shock (*p* = 0.010). Therefore, the direction of this effect suggests that male mice may be more sensitive to lower shock intensities, which is not sufficient to explain the overall decrease in responding we see in the punished responding paradigm and if anything suggests that females are even more sensitive to the behavioral effects in the absence of enhanced shock sensitivity.

**Figure 5.**
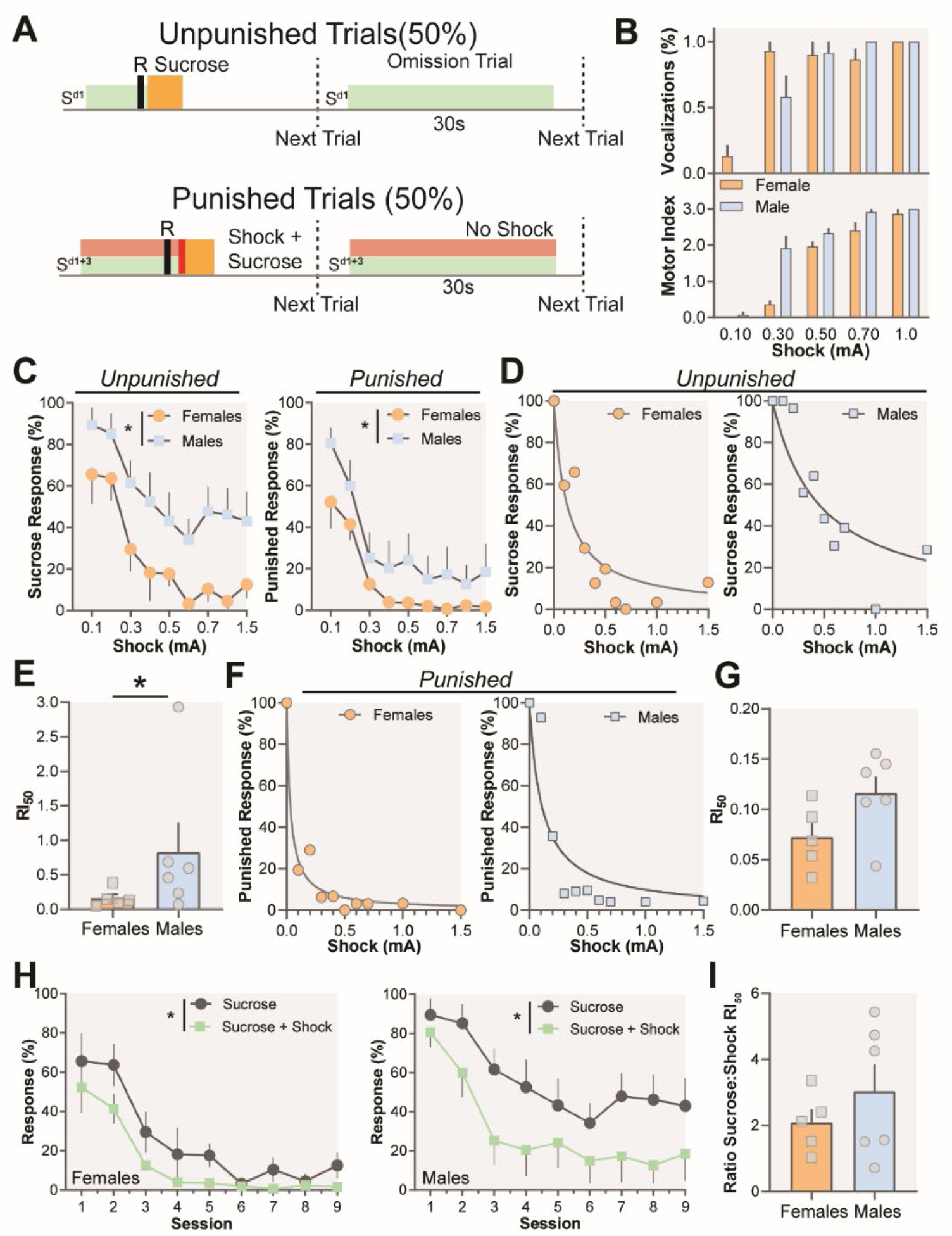
Females are more sensitive to punishment. (A) Schematic of punished responding. (B) Males were more sensitive to unsignaled shocks of varying intensities. There is a sex-based difference in motor response (Sex x Shock Intensity Interaction, F(4,35)=2.802, p=0.0406; Sex Main Effect, F(1,35)= 1.830, p>0.05; Shock Intensity Main Effect, F(4,35)= 48.33, p<0.0001) and vocalizations (Sex x Shock Intensity Interaction, F(4,35)=6.991, p=0.0003; Sex Main Effect, F(1,35)= 27.58, p<0.0001; Shock Intensity Main Effect, F(4,35)= 111.3, p<0.0001) with increasing shock intensities. (C) Males respond more during both unpunished (left) and punished (right) trials as compared to females over increasing shock intensities (Sex x Shock Intensity Interaction, F(8, 72) = 0.3219, p=0.9552; Sex Main Effect, F(1, 9) = 5.157, p=0.0493; Shock Intensity Main Effect, F(2.930, 26.37) = 15.16, p<0.0001) and punished responses (Sex x Shock Intensity Interaction, F(8, 72) = 1.259, p>0.05; Sex Main Effect, F(1, 9) = 4.644, p=0.0596; Shock Intensity Main Effect, F(2.074, 18.67) = 41.02, p<0.0001). (D) Representative response inhibition 50 curves (RI50) for sucrose responding for males and females. (E) Males show a higher RI50 than females for sucrose responding (Mann-Whitney U=5, p=0.0480) indicating that more shock intensity was necessary to reduce their response rates. (F) Representative response inhibition 50 curves (RI50) for punished responding for males and females. (G) There was a trend towards higher RI50 in males during punishment (t(9)= 1.971, p=0.1810). (H) In comparing trial type, there is a main effect for males (F(1, 10) = 10.90, p=0.0080) but not for females (F(1, 8) = 3.812, p=0.0867). (I) RI50 (sucrose responding RI50/punished responding RI50) is not different between sexes (t(9)=0.9748, p=0.3551).

### The MCOAT highlights the complexities of sexually dimorphic behaviors (Fig 6)

Together these data highlight sex-specific behavioral strategies that guide context-specific behavioral performance. While females self-administer greater levels of sucrose they acquire negative reinforcement at slower rates. However, once acquired, females’ behavioral bias is towards avoiding aversive outcomes, rather than seeking rewards. Males have different strategies that are more biased towards maximizing rewarding outcomes - however, in situations where aversive outcomes are certain their behavior shifts to avoiding them. Together, these differences are the foundation for learned behavior and are important substrates to consider in expression and development of disease etiology.

**Figure 6.**
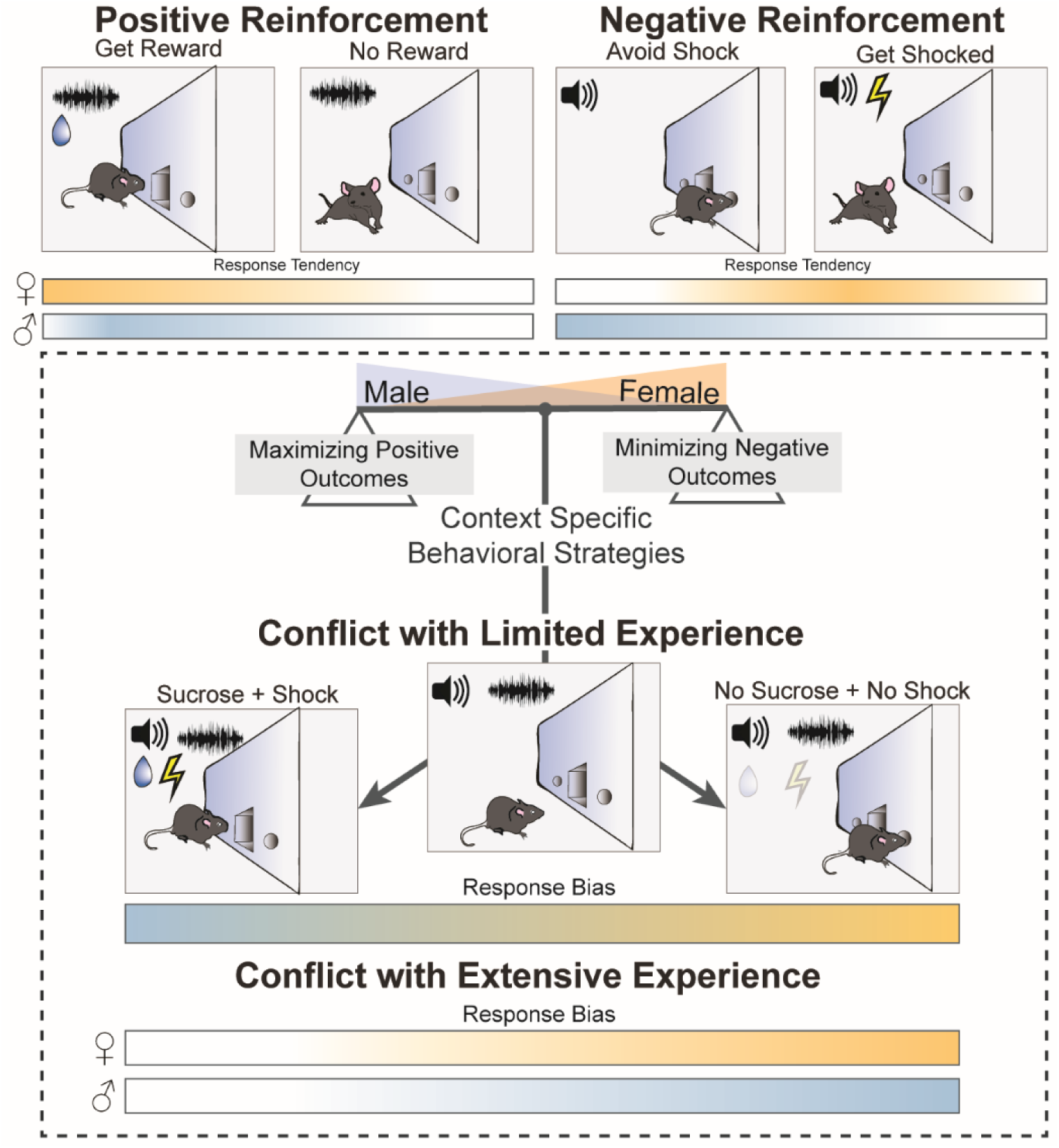
Females are biased towards avoiding aversive outcomes. Males and females demonstrate divergent rates of learning for positive and reinforcement; however, while females learn negative reinforcement at a slower rate they are biased towards avoiding aversive outcomes when they are presented concurrently with options to obtain rewards. Further, females demonstrate a bias for punishment avoidance and males demonstrate a bias for reward-seeking in the face of an aversive stimulus.

## Discussion

Emerging evidence has highlighted that biological sex itself is not a behavioral determinant, but rather is a complex variable interacting with environmental factors and experience to drive behavior, thus requiring studies that allow for an understanding of the behavioral factors that underlie sex-specific strategies (18–21). Here we present and validate a complex behavioral task (Multidimensional Cue Outcome Action Task, MCOAT) that can dissociate a variety of factors that interact to drive goal-directed behavior. We then use this task to demonstrate sex differences in fundamental behavioral strategies that manifest in the execution of learned behavior. First, we showed that female mice self-administer higher levels of sucrose, but acquire negative reinforcement at a slower rate; however, in situations where positive and negative stimuli are presented together, females favor avoiding aversive outcomes over seeking rewards. To further support the hypothesis that females are more motivated by the avoidance of aversive outcomes, we show that in situations where a cued shock is delivered concurrently with a sucrose delivery in a percentage of trials females reduce responding at lower shock intensities. Additionally, females reduced responding over all trial types in the task - regardless of whether the punisher was presented in those trials or not. Together, fundamental differences in basic behavioral strategies between the sexes - specifically in regard to the balance of positive and negative outcomes - point to critical factors that may guide behavior in females and may underlie the differences in development and progression of psychiatric disease states (21). These findings are particularly important as they highlight the dangers of making generalizations about female-specific behavioral strategies with unidimensional behavioral tasks that present stimuli in isolation.

Decision-making is a process in which various external and internal processes are in play to ensure homeostasis between positive and negative outcomes (22). This balance allows organisms to assess risks of aversive outcomes while obtaining potential rewards. Here we present evidence demonstrating that female mice have an intrinsic bias towards avoiding negative outcomes over obtaining rewards. On the other hand, male mice showed a similar bias only when they had extensive experience with cue-outcome contingencies suggesting intrinsically male mice may make riskier decisions by seeking out rewards without considering aversive outcomes to the same extent. Importantly, the behavioral bias that female mice showed was not due to a differential ability to perform the task, as at the end of the negative reinforcement and discrimination phases, both sexes exhibited similar levels of performance. Therefore, although the behavioral bias for avoiding aversive outcomes is affected by experience, at the basal level it shows a strong sex difference, which may have evolutionary basis. In mammals, parental behavior has evolved to be sexually dimorphic (23). That is, while males usually engage in indirect parental investment (e.g., foraging), females perform direct parenting (e.g., nursing, rearing offspring, shelter construction), which requires them to be in close proximity to their offspring (23). The high level of direct maternal investment including strong dependence of the offspring on the mother for its survival has been shown to be the main limiting factor for female risk-taking behavior (24, 25). Therefore, there is strong sexual selection pressure for female mammals to be risk-averse and value avoidance of aversive outcomes highly. This evolutionary mechanism can explain our findings showing that female mice biased to choose avoiding footshocks over a sucrose reward even in the conditions where stimulus-outcome contingencies were not completely established. It is important to note, however, environmental factors and learning influence this behavior in males’ risk-taking strategy indicating a plastic rather than a fully deterministic process.

Our results highlight that stimulus specific learning is an important factor in the expression of sex differences and shows the importance of understanding context-specific behavior when making conclusions about sex-specific strategies. We show that sex differences in learning rates are dependent on stimulus value whereby females learn behavior reinforced by a negative stimulus (shock) at a slower rate as compared to males, while learning rates are the same when reinforced by a positive stimulus (sucrose). Importantly, during this task, the motoric action associated with both of the stimuli are identical (nose-poke response), allowing us to definitively determine that these effects are not driven by differences in movement. Similarly, correct responses on both trial types results in a positive outcome (delivery of sucrose, or removal of shock) which allows for specific determination that these effects are driven by the stimulus value rather than the outcome value. Indeed, the ability to look in the same experimental subject over tasks with divergent stimuli (sucrose vs. footshock) but convergent (nose-poke response) outcomes OR convergent stimuli (shock) with divergent behavioral outcomes (i.e. response vs response inhibition) is a major advantage of the MCOAT. In the initial phase of the task, we show that females self-administer more sucrose as compared to males, which is in line with previous studies reporting increased reward seeking in females for both natural (26, 27) and drug reinforcers (10, 28–30). These previous findings have led to the conclusion that females are more driven by positive outcomes. However, females also performed worse on the negative reinforcement task as well as in the discrimination phase, suggesting stimulus specific processing, not “reward” per se as the removal of a shock is a positive outcome. This strongly suggests that simple behavioral tasks developed to test a single dimension of behavior may lead to incomplete conclusions about sex differences.

While we used the MCOAT within this study to elucidate sex-specific behavioral strategies, the power of this task is its potential for application across the mental health fields. This behavioral procedure will be particularly powerful for examining the effects of many other external (e.g., stress, depression, addiction) or internal (hunger, thirst) conditions on learned behavior, ongoing performance, behavioral biases, and goal-directed action. The MCOAT - and similar behavioral tasks - address a recent focus of the National Institutes of Health (NIH) to utilize and develop complex behavioral models in order to increase the translatability of findings. These findings also further highlight the importance of this initiative where conclusions drawn from simple tasks conflict with the conclusions that are able to be drawn from complex tasks across a number of modalities (20). This is highlighted clearly with the *Phase 1* data showing increased sucrose consumption in females and decreased rates of negative reinforcement learning, which would lead to the conclusion that females are more reward driven, when in fact, the comprehensive data suggests the exact opposite. Understanding these processes precisely is critically important to improving treatments for these conditions, especially in women where treatment efficacy is reduced and off-target and adverse consequences from medications are particularly high (31–33).

While the MCOAT will be a valuable tool for a number of subfields within the greater behavioral neuroscience discipline, its application may be even more powerful when paired with neural recording and manipulation techniques. One of the difficulties with defining the precise information that is encoded within cellular populations in the brain is the ability to dissociate the conflating factors that contribute to behavior in standard tasks that were not designed for this purpose. During a behavioral response salience, novelty, value, prediction, action initiation/motor responses, sensory information, among other things occur simultaneously and can contribute to temporally-specific neuronal signatures occurring around these discrete behavioral events. Further, learning-theory models take into account many of these factors and predict changes in learning rate and performance if any one of them is altered (34–36). Thus, manipulation techniques that alter only one of these factors (salience, but not value, for example) will alter learning rates in a manner that would be predicted based on the other factor that is intended to be tested. Most of the current tasks that are used in the neural encoding field rely on simple Pavlovian schedules where animals are presented aversive or rewarding stimuli. In most of these tasks, the cue presentation and the conditioned response occur simultaneously, meaning that an increase in neural activity to the cue could encode a response, prediction, salience, attention, or cue value. Similarly, in simple operant tasks where animals lever press for a single stimulus without competing environmental information the signal could be motor activity, outcome value, prediction, etc. Thus, understanding how cellular populations in the brain encode information relies entirely on our understanding of the complexities of behavioral responses. The MCOAT disentangles the specific encoding of these stimuli and is designed to independently probe stimulus value, outcome, cue learning, and action, because the discrete phases allow for these factors to be manipulated independent of one another. For example, punishment and negative reinforcement have the same outcome - avoidance of shock - but opposite behaviors - nose-poke or inhibit nose-poke - so if a circuit encodes the action it will be divergently activated during each discrete phase and if it encodes the outcome it will be similarly activated. Together the MCOAT allows for a dissociation of neural responses to cues, outcomes, actions, and value-based stimuli to dissociate and isolate these critical aspects of behavioral control.

Overall, the studies we report here highlight the potential pitfalls of making general conclusions based on behavioral procedures using a single stimulus and/or contingency. We also demonstrate the importance of studying reinforcement learning across multiple environmental conditions in order to establish how sex differences are expressed in preclinical and clinical models. As mentioned above, it is imperative to consider that differences in fundamental behavioral strategies that can interact with environmental conditions to drive behavior. Finally, we developed a novel behavioral task (MCOAT) to achieve this critical behavioral feature and successfully employed this task to show that female mice are biased towards avoiding aversive outcomes during conflict even though they learned avoidance behavior at a slower rate. Together, we present a set of experiments defining the sex-specific behavioral strategies that guide value-based decisions in males and females, and show the underlying the complexity and importance of understanding sex differences using composite behavioral tools.

## Acknowledgements

This work was supported by NIH grants DA042111, DA048931 to E.S.C., GM07628 to J.E.Z., MH065215 and DA047777 to A.R.J, MH064913-15 to K.C.T., and K00DA048436 to A.J.L. as well as by funds from the VUMC Faculty Research Scholar Award to M.G.K., the Vanderbilt Academic Pathways Fellowship to L.J.B., Brain and Behavior Research Foundation to E.S.C., the Whitehall Foundation to E.S.C., and the Edward Mallincrodt Jr. Foundation to E.S.C.

## Disclosures

We declare no conflict of interests.

---

**Supplementary Figure 1.**
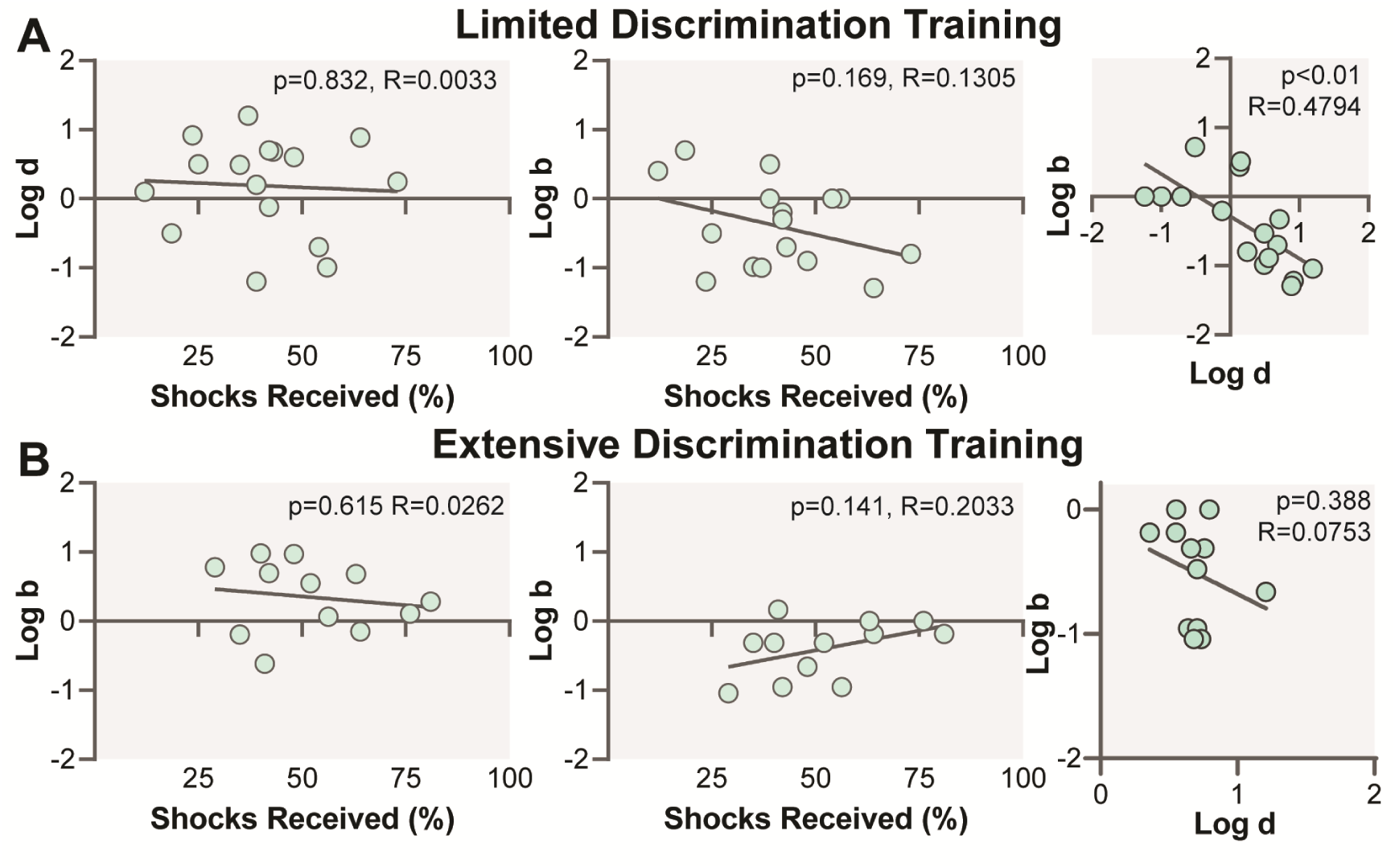
**(A)** Percent of total shocks received over the course of negative reinforcement training does not correlate with Log *d* (*p*>0.05, R=0.0033) or Log *b* (*p*>0.05, R=0.1305) in limited discrimination training or in **(B)** extensive discrimination training (Log *d, p*>0.05, R=0.0262) (Log *b, p*>0.05, R=0.2033). **(A)** There is a significant negative correlation between Log *b* and Log *d* in limited discrimination (*p*<0.01, R=0.4794) and **(B)** extensive discrimination (p<0.05, R=0.0753).

**Supplementary Figure 2.**
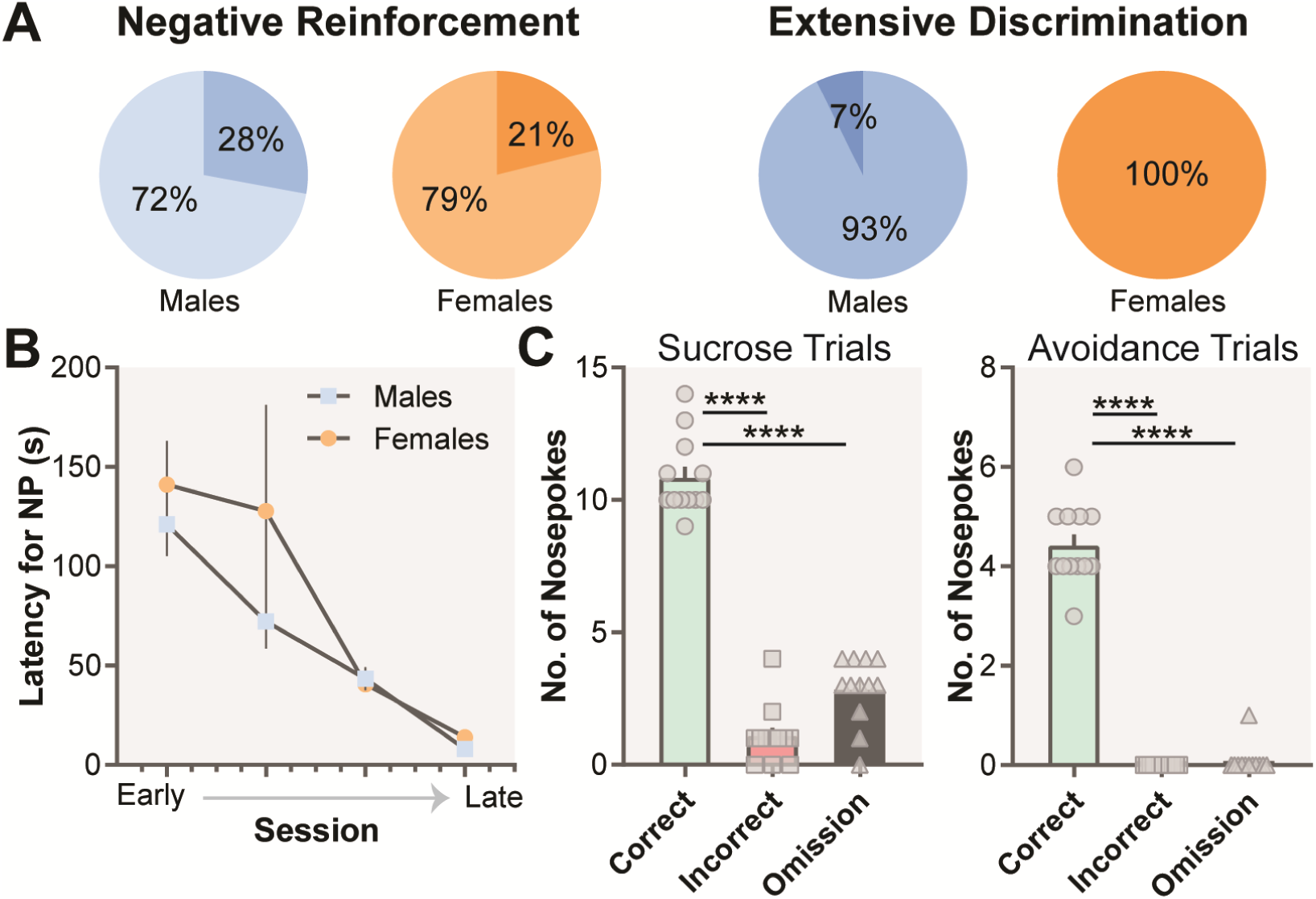
**(A)** In lightest shade, the percentage of mice that escaped shocks. In medium shade, the percentage of mice that avoided shocks. In darkest shade, the percentage of mice that omitted trials. **(B)** Latency to nose-poke following cue-onset (S^d2^) decreases with training during negative reinforcement (*p>*0.05). **(C)** At the end of extensive discrimination, animals make a high number of responses for sucrose with comparatively few incorrect (p<0.0001) and omitted (p<0.0001) responses. Animals respond similarly at the active nose-poke to avoid/escape shocks and make comparatively few incorrect (p<0.0001) and omitted (p<0.0001) responses.

**Supplementary Figure 3.**
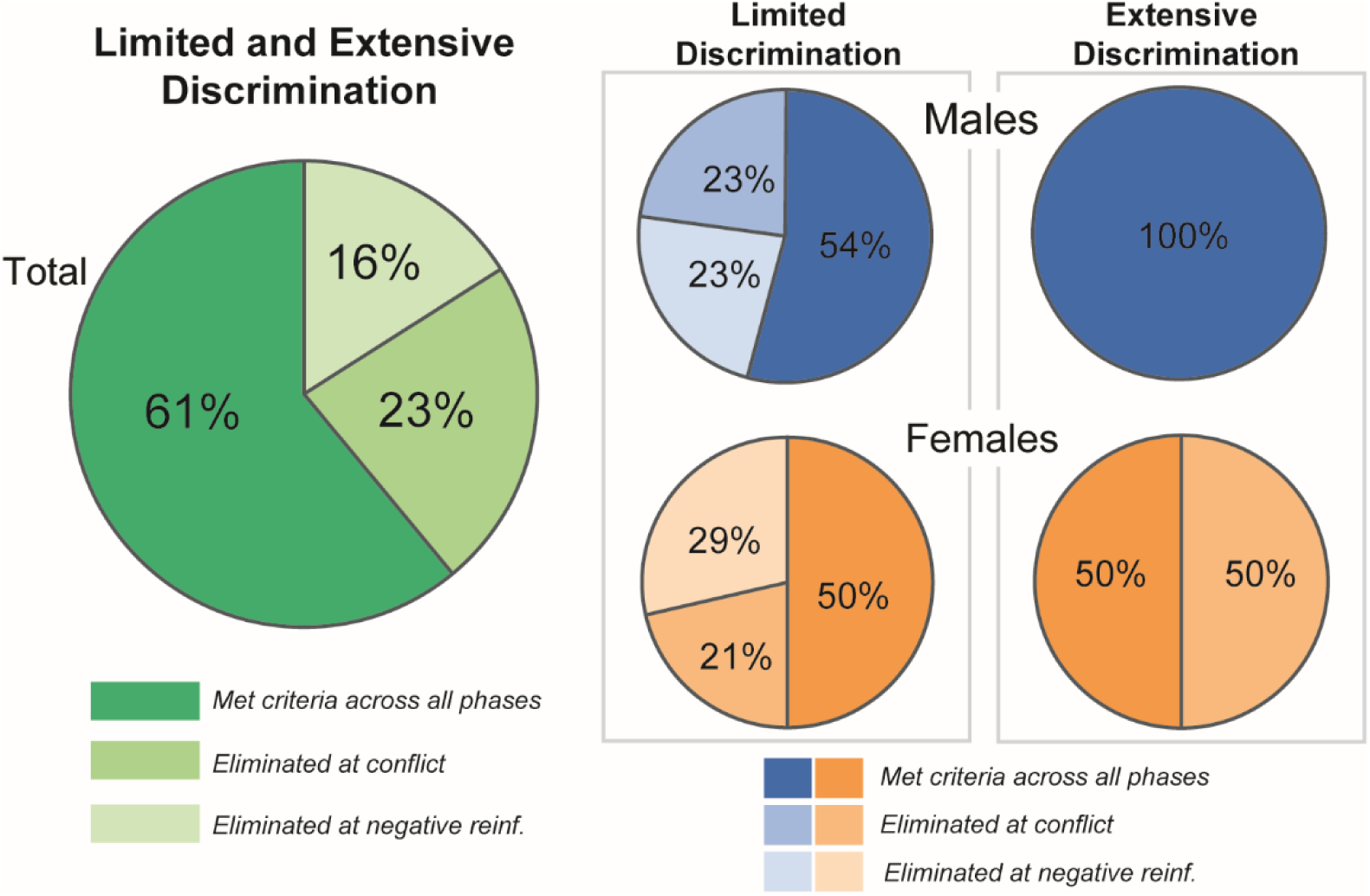
Total mice that were eliminated from the study during negative reinforcement, during conflict, and remaining that met criteria across all phases limited and extensive discrimination combined. Fifty four percent of males and 50% females completed all phases of limited discrimination training. One hundred percent of males completed Phase 1 and Phase 3 of extensive discrimination. Fifty percent of the females failed to meet criterion and 50% completed both phases with extensive training.

